# Responses to taste and eating in the parabrachial nucleus of the pons in awake, unrestrained rats

**DOI:** 10.1101/2025.08.30.673255

**Authors:** Flynn P. O’Connell, Joshua D. Sammons, Steven A. Pilato, Patricia M. Di Lorenzo

## Abstract

**Background/Objectives.:** The parabrachial nucleus of the pons (PbN) is a hub in the central pathway for taste in non-primate mammals. Recent evidence has identified a role for the PbN in regulating ingestion; however, little is known about how the PbN responds to solid food. Here, we recorded PbN responses to liquid taste stimuli over days/weeks, and tested whether these responses are good predictors of responses to solid foods consumed in a naturalistic way.

**Methods.:** Rats were prepared for one-photon calcium imaging by surgical implantation of a GRIN lens. PbN activity was imaged during multiple sessions spanning up to 112 days while animals licked various tastants in an experimental chamber (Lick phase). In some sessions, following the Lick phase, rats were presented with Granny Smith apples, milk chocolate and/or salted peanuts (Food phase). In one session, chocolate or peanut odor was presented during the Lick phase along with the tastants.

**Results.:** PbN cells responded to more than one taste quality with tastant-specific spatiotemporal patterns of response. While response profiles of individual PbN cells shifted over days/weeks, across-unit patterns differentiating taste quality remained relatively stable. Population responses to liquid tastants and solid foods were segregated according to the different motor patterns required for ingestion. Responses to liquid tastants were not good predictors of responses to solid foods. Importantly, PbN cells also responded to food-related odorants.

**Conclusions.:** Collectively, these results challenge the traditional view of the PbN as a simple relay for taste and instead position it as an integrative hub involved in processing gustatory, olfactory, and somatosensory information. Moreover, the findings emphasize the importance of population coding in maintaining perceptual stability.

## Introduction

The medial parabrachial nucleus of the pons (PbN) has been characterized as a “taste relay” because it is an obligatory synapse between the taste-related portion of the nucleus tractus solitarius (NTS) and more rostral structures in the gustatory pathway [1] in non-primate mammals. In addition to processing information about taste, recent evidence has positioned the PbN squarely in the neural circuit that regulates ingestion [2–5]. Although there is abundant information about PbN cells’ responses to liquid taste stimuli, there is a little evidence about how cells in the PbN respond during ingestion of solid food. In fact, the great majority of studies of taste in the PbN have been conducted in anesthetized animals, obviously precluding study of responses during consumption. Moreover, rather than food-related stimuli, exemplars of the “basic” taste qualities (sweet, salty, sour, bitter and umami) have been used most commonly as stimuli to probe the system’s functionality. When food-related, naturalistic tastants were used as stimuli [6], results suggested that characteristics in addition to taste, such as odor and texture, may collaborate to convey information about food in the PbN. Thus, multimodal responses evoked by solid food may be crucial for decisions about consumption.

Most studies of PbN taste responses are the product of recording from one cell at a time, with the aggregate used to deduce theories of taste coding. For example, for each taste-responsive neuron, the relative response rates evoked by prototypical tastants representing the basic taste qualities defines its “tuning profile”. Narrowness or broadness of tuning across this stimulus array have served as the basis of theories of the general coding scheme used by the population. Historically, two theories have dominated the literature: the labeled line and the across unit pattern theories (reviewed in [7]). According to the labeled line theory in its most extreme form, each taste quality is represented by nonoverlapping groups of cells that respond “best” to exemplars of that quality. Responses to tastants other than the best stimulus are thought to be “sideband” responses that carry largely irrelevant information. An alternative conceptualization is the across-unit pattern theory. This theory is based on the observation that even though the response to one taste quality may dominate, most taste cells respond to multiple taste qualities. This theory argues that each stimulus evokes a unique pattern of responses across the population. Population response patterns associated with similar tastes are highly correlated, while patterns evoked by stimuli of different taste qualities are poorly correlated.

For the labeled theory in particular, an inherent assumption is the stability of neuronal tuning profiles over time. So, for example, a NaCl-best cell will always be a NaCl-best cell and its inputs and outputs will form an information channel related to NaCl. Similar logic is applied to the other taste qualities leading to the idea that there are five such information channels related to the five basic qualities. The across unit pattern theory, on the other hand, does not require the assumption of stability of individual tuning profiles over time, only that the patterns evoked by the various taste qualities remain identifiable and different from those evoked by other taste qualities. In the present study, we tested the hypothesis that tuning profiles of PbN taste-responsive cells remain a stable characteristic over time.

Although much insight can be obtained from recording responses to basic taste qualities, to fully assess the role of the PbN in appetitive and consummatory behaviors it is necessary to record how the system functions when the animal is actually eating food. In that regard, investigations of responses to tastants that are licked and swallowed have provided some insight into how the taste of food might be encoded in the PbN. For example, in studies of PbN [8] and NTS [9] taste-responsive cells recorded in awake, unrestrained rats, a substantial subset of cells showed long latency (>1.5 s) responses to taste stimuli acquired through a rapid (<∼1 s) sequence of licks. This type of response suggests that stimulation of extraoral taste receptors might contribute to the brainstem response to taste/food. In addition, Sammons et al. [6] showed that naturalistic taste stimuli (e.g., grape juice, clam juice, lemon juice, coffee) produced more discriminable responses than responses to prototypical tastants (e.g., sucrose, NaCl, citric acid, quinine, monosodium glutamate) across the population of PbN cells. These data suggest that food, with its varied sensory components including taste, odor and texture, might be more effective in driving the system compared with the prototypical taste stimuli. Finally, taste-responsive cells in both PbN [8] and NTS [9] are intermingled with cells that showed lick-related activity, suggesting that the act of food consumption writ large is reflected in the cellular activity in these brainstem areas.

In a recent study of the NTS by Pilato et al. [10], electrophysiological responses to licking prototypical tastants and to eating solid foods were recorded. Foods were chosen to emulate each of the basic taste qualities. Specifically, the foods were milk chocolate (sweet), salted peanuts (salty), Granny Smith apples (sour), and broccoli (bitter). Results showed that responses to licking sucrose, NaCl, citric acid and quinine were not predictive of responses to the foods matched for the dominant taste quality. The most robust responses from NTS cells were recorded during the appetitive phase of ingestion when the rat’s head was approaching and exploring food in the food well. It is possible that the animal’s tongue was in motion in anticipation of contact with the food so that the responses prior to contact with the food reflected movement rather than a cognitive state of anticipation. Another possibility is that these early responses might be food-related olfactory responses. In contrast, while the animal was actually ingesting the food, a large portion of NTS cells became quiescent.

Here, using one-photon calcium imaging of the PbN, we explored the two aspects of the neural representation of taste and food discussed above. First, we examined the stability of taste responses in PbN cells over time, i.e., several days or months. Results suggest that PbN cells change their tuning profiles over time, but the representation of the various taste qualities remains discriminable through population coding. Second, we compared the PbN responses to licking liquid tastants and eating solid food. Results revealed that licking tastants evoked taste-specific spatiotemporal patterns of response, with most cells being muti-sensitive across tastants. In addition, we found that cells in the PbN respond to food-related odors. In contrast, responses to solid foods were more closely tied to the motor components associated with appetitive and consummatory behaviors. Finally, as in the NTS [10], responses to liquid tastants were not predictive of the responses to solid foods containing a congruent taste component.

## Materials and Methods

### 2.1 Subjects

Four female Sprague Dawley rats (Taconic Labs) weighing 250–350 g served as subjects for these experiments. Animals were maintained on a 12:12-h light-dark cycle (lights off at 0900 hours) and pair-housed in plastic cages with environmental enrichment until surgery. Following surgery, the animals were single housed. Standard chow and water were available *ad libitum*. During data collection, animals received one hour of water daily in addition to fluid consumed during the experimental paradigm. Because these animals were run with intervals spanning days to weeks, they were water-deprived and lick trained (see below) for several days prior to image recording. They were then returned to *ad libitum* water. All procedures were approved by the Binghamton University Institutional Animal Care and Use Committee.

### 2.2 Implantation of a Gradient Refractive Index (GRIN) lens for calcium imaging

Approximately 15-20 min prior to initiating anesthesia, subjects were administered buprenorphine (0.01 mg/kg, s.c.) and atropine (0.05 mg/kg, s.c.). Each rat was then placed in an induction chamber and exposed to 3% isoflurane for 10 min. Throughout the surgery, anesthesia was maintained at 1-3% isoflurane, with adjustments made as necessary. The rat’s scalp was shaved and its head secured in a stereotaxic instrument (David Kopf Instruments, Model 1900, Tujunga, CA) using blunt ear bars. The head was positioned downward at a 25° angle. Artificial tear gel was applied to the eyes to prevent drying, and a rectal thermometer connected to a temperature regulator and heating pad maintained body temperature at 37° C. The scalp was cleansed alternately with Betadine and 70% ethanol three times. After drying, a midline incision was made in the scalp, and the fascia were retracted through blunt dissection. A hole was drilled above the PbN (−11.3 A/P from bregma, 1.7 M/L). The exposed dura was removed and a Hamilton syringe was lowered ∼5.6 mm below the surface of the brain into the PbN to deliver 0.5 µL per hemisphere of an AAV9 virus containing the GCaMP7s gene under the synapsin 1 promoter (Addgene #104487-AAV9; pGP-AAV-syn-jGCaMP7s-WPRE). The wound was then closed with nylon suture and treated with Neosporin ointment. Anesthesia was then stopped. The rat was then placed in a holding cage until it regained consciousness and mobility after which it was returned to its home cage. Subsequently, 0.05 mg of buprenorphine, 3 mL of warmed saline, and 0.05 mg gentamicin were administered s.c. Buprenorphine and gentamicin were administered daily for three days post-surgery. The rats were weighed daily until they reached 90% of their pre-surgical body weight.

Following two to four weeks of recovery, the rats were re-anesthetized, as above, and six stainless steel skull screws were embedded in the skull. A GRIN lens (10 mm long; 1 mm diameter; Inscopix, Inc., Mountain View, CA) attached to a baseplate was implanted in the brain such that the ventral tip was positioned just above the PbN. The baseplate and GRIN lens were secured to the skull and skull screws with dental acrylic. The wound was then closed with nylon suture and treated with Neosporin ointment. Anesthesia was then terminated and postoperative care was administered as described above.

### 2.3 Apparatus

Data collection was conducted within a transparent Plexiglas experimental chamber (Med Associates, Fairfax, VT) enclosed in a melamine box. A clear Plexiglas window on the front door of the melamine box facilitated observation and video recording of the rat’s behavior. Positioned just behind an aperture on one wall of the experimental chamber was a stainless steel sipper tube. Lick events were registered when the animal interrupted an infrared photobeam directed across the aperture. The sipper tube contained an array of smaller stainless steel tubes, each connected to a dedicated pressurized reservoir of a taste stimulus. A computer-triggered solenoid regulated the flow of fluid from the tastant reservoir to the sipper tube. Upon each lick, excluding “dry licks,” a volume of 12 µl ± 2 µl of fluid was dispensed within 10 ms. This arrangement enabled precise control over the stimulus administered to the rat on a lick-by-lick basis. The fluid delivery system was calibrated daily. “Food wells,” consisting of stainless steel boxes, 5 cm X 5 cm X 15 cm, open to the inside of the experimental chamber, were located at its four corners (see **Fig 1B**). An infrared photobeam across the opening of the food wells was used to record the time that the rat’s head breached the opening. The food wells were blocked by opaque panels during the “Lick phase” of the experimental procedure, after which the panels were removed, and the food wells were then filled with solid food for the “Food phase” of the experiment.

**Figure 1.**
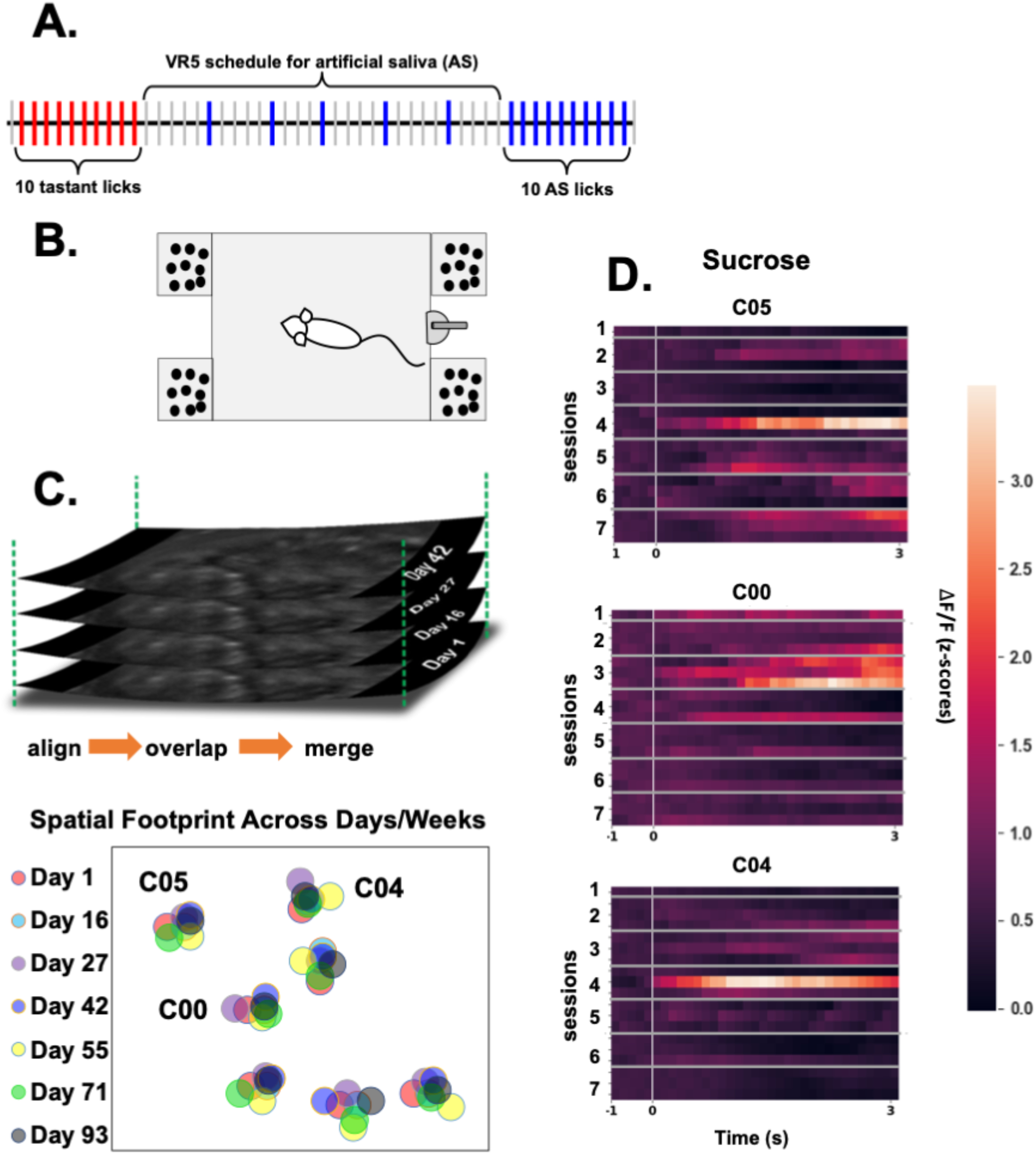
**A**. schematic illustrating the tastant presentation method for the Lick phase. Ten consecutive tastant licks were preceded and followed by five licks of artificial saliva (AS) presented on a variable ratio five schedule (VR5). That is, AS licks were separated by 4 to 6 unreinforced (dry) licks. The panel also illustrates that AS was also presented as a stimulus. Tastants and AS trials occurred in random order. **B.** Schematic showing the arrangement of food wells at the corners of the experimental chamber, along with a drinking spout between the two food wells on the righthand side. **C.** Schematic illustrating the method for determining the presence of the same cells over different calcium imaging sessions. Lower panel shows the results of this analysis. Seven cells were present in the same location across sessions for 93 days in animal PGT08. **D.** Heatmaps showing the responses to licking sucrose across seven sessions in three different cells in animal PGT08. In all three cells, responses were highly variable across trials within a session and across days.

### 2.4 Taste, food and olfactory stimuli

Taste stimuli were exemplars of the five basic taste qualities: 0.1 M sucrose (S) for sweet, 0.1 M NaCl (N) for salty, 0.0167 M citric acid (CA) for sour, 0.0001 M quinine HCl (Q) for bitter, and 0.1 M monosodium glutamate (MSG) plus inosine monophosphate (0.01 M) for umami. Tastants were made with reagent grade chemicals (Fisher Scientific, Pittsburgh), and dissolved in artificial saliva (AS; 0.015 M NaCl, 0.022 M KCl, 0.003 M CaCl_2_ and 0.0006 M MgCl_2_ at a pH of 5.8 ± 0.2; [11]. AS was also used as a taste stimulus.

In some experimental sessions, two olfactory stimuli were presented during the Lick phase of the experimental procedure. These consisted of chocolate and peanut odors. Milk chocolate bits and chopped peanuts were loaded into separate bottles fitted with tubes for delivery of compressed air and expulsion of odiferous air. Odorants were conveyed to an olfactory port adjacent to the lick spout in the experimental chamber. During the odorant presentations, the rat was licking artificial saliva.

Food stimuli were selected based on the dominant taste quality that they conveyed. They were: Nestle milk chocolate chips (35% cacao, 67% sugar; sweet), dry roasted salted nuts (salty), and Granny Smith apples (sour). In some sessions, only chocolate and peanuts were offered to complement the olfactory stimuli. Three gms of each food were loaded into the food wells after the Lick phase concluded.

### 2.5. Experimental procedures

#### Lick Training

Following recovery from surgery, rats were water deprived for 18-23 hr and trained to lick sucrose from a lick spout in the experimental chamber. This training regimen involved daily sessions lasting 30 min (excluding weekends), during which 0.1 M sucrose dissolved in AS was presented at the lick spout. Imaging sessions began once the rat consistently achieved > 500 licks per day for three consecutive days. Water deprivation was maintained throughout the imaging sessions and paused on weekends.

#### Experimental Sessions

Each experimental session consisted of two 30 min phases: the Lick phase and the Food phase. The Lick phase always preceded the Food phase. During the Lick phase, rats were provided access to the sipper tube, which dispensed 12 µL ± 2 µL per lick of tastant as described above. Each taste trial consisted of 10 consecutive licks of a tastant, followed and preceded by six licks delivered on a variable ratio five schedule (see **Fig. 1A**). That is, four to six dry licks were interspersed with licks of AS. AS also served as a “taste” stimulus. In addition, two odorants were delivered through an odor port directly adjacent to the sipper tube. Rats licked AS for 10 licks during the 2 s odor trial. In the Food phase, which followed immediately after the Lick phase, rats were allowed to explore and consume solid foods placed in the food wells.

#### Calcium imaging

A miniscope used for calcium imaging (Inscopix, Inc., Mountain View, CA) was fixed to the baseplate on the animal’s head prior to the animal being placed in the experimental chamber. The depth of the GRIN lens was adjusted to maximize the number of cells in the field that could be detected. Animals were run for multiple sessions; the number of cells detected in each session varied. Some cells were only detected in response to specific tastants.

### 2.6. Data Analyses

Raw videos were preprocessed in Mosaic software (Inscopix, Inc., version 1.8.1). Spatial downsampling (2× binning) was applied to reduce data volume and computational overhead. Motion correction was performed using a rigid body registration algorithm that aligned all frames to the initial frame of the session. In some cases, motion correction was applied iteratively to improve alignment. After registration, frames were spatially cropped to exclude regions out of focus.

Regions of interest (ROIs) corresponding to putative neurons were initially extracted using Inscopix’s automated segmentation pipeline based on constrained nonnegative matrix factorization (CNMFe). ROIs were manually refined, when necessary, with cell masks examined by one of the authors to confirm neuron morphology and spatial separation from neighboring cells. Extracted calcium traces were visually inspected to remove any non-neuronal signals or imaging artifacts. Prior to visualization and downstream analyses, calcium traces were z-scored, and negative z-scored values (below baseline) were set to zero.

To facilitate longitudinal analyses of neuronal activity, neurons were retrospectively matched across imaging sessions spanning several months. Given the relatively low neuronal density, neurons could be reliably identified using summary representations of neuronal activity, such as mean-subtracted images and ΔF/F movies normalized to the mean activity of the entire recording. Anatomical landmarks, particularly blood vessels, further guided neuron identification. Neurons consistently identified across sessions were assigned unique indices and grouped for downstream analysis.

Event timestamps were derived from synchronized GPIO channels and aligned to the imaging time base using a nearest-match algorithm. Lick-related and stimulus-related timestamps were used to define trials and separate reinforced from non-reinforced activity periods. For each trial, a 4-s pre-stimulus baseline was computed and used to normalize data to ΔF/F. The response window was defined as a 2-s interval centered at the peak of the calcium transient following stimulus onset. If overlap occurred between the baseline and response windows, the response window was shifted to begin at the first stimulus lick. The full analysis window extended up to 5 seconds after the final lick. A response was considered significant if the mean signal during the response window exceeded 2.58 standard deviations above the mean baseline.

The breadth of tuning was determined by calculation of Uncertainty (*H*, [12]), an index of the relative magnitudes of response across tastants. This measure is calculated as follows:

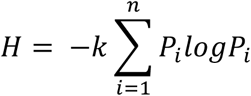

where *n* is the number of tastants (5), *k* = 1.4307 for 5 tastants, and *P_i_* is the ratio of tastant *i* response magnitude to the sum of all tastant response magnitudes. The value ranges from 0, signifying that the neuron responds to a single tastant, to 1.0 indicating that the neuron responds to all 5 tastants equally.

For a subset of sessions containing eating episodes, event intervals were defined based on the animal’s head position: specifically, when the animal entered the food well to retrieve food, exited the well to return to its designated eating spot, and subsequently began eating. Using multiple synchronized camera views, researchers manually annotated these distinct behaviors as eating events for each food type, as well as other behavioral states including quiescence and grooming.

#### 2.6.1. Principal Component Analysis (PCA)

To further characterize taste-related patterns in neuronal activity, PCA was performed on taste-responsive neurons (defined as neurons exhibiting activity ≥ 2.58 SD above baseline). Initially, neuronal activity was averaged across trials for each tastant to generate trial-averaged responses prior to PCA (Fig. 3, PGT13). Complementary analyses were also conducted on the full dataset without trial averaging: first, by collapsing all tastants into a single “Licking” event without eating (Fig. 8D, PGT08), and second, with licking events including eating (Fig. 9B, PGT13). Last, PCA was performed without trial averaging, labeling each individual trial separately. This analysis demonstrated clear separation among primary behavioral events such as grooming, eating, tastant exposure, and odorant exposure (Fig. 12B, PGT13).

For Fig. 7D, to view possible state transitions among the 12 PbN cells recorded during the session, dimensionality reduction was performed using PCA on neural activity that occurred during a subset of licking and no licking intervals (500 seconds in total). A time by neuron rectangular matrix was created before implementing the dimensionality reduction in R statistical software. The first three principal components were used for visualizing population time courses. The time series were grouped by licking behavior in order to assess transitions that occur as the animal’s behavior changed.

For Fig. 9A, dimensionality reduction was performed on the neural activity of a population of 6 PbN neurons, while the rat ate solid food. The time courses included grooming, well entry (food wells contained nuts, milk chocolate, or apple), and eating (eating apple).

## Results

Responses to taste stimuli were imaged from 56 cells in the PbN of four awake, unrestrained rats (PGT06, 08, 09, 13). In two rats (PGT09, 13), responses to eating solid foods were also imaged and in one rat (PGT13), responses to food-related odors were imaged along with responses to tastants and solid foods. In three rats, 29 cells were identified that were present across sessions. Specifically, in rat PGT06, 11 cells were present in two sessions spanning seven days.; in PGT08, seven cells were present in seven sessions over 99 days; in PGT13, 11 cells were present in eight sessions over 112 days.

### Taste response profiles over time

**Fig. 1C** illustrates the method used to verify the consistent presence of seven cells across seven sessions spanning 99 days in rat PGT08. Images from each session were aligned and units present in all sessions were identified (see Materials and Methods). In each of these sessions, other cells were present, but were not consistently present over repeated sessions. This was true for all three rats where cells were imaged over time.

In general, taste-responsive cells changed their responsiveness from session to session and even from trial to trial. **Fig. 1D** illustrates this point. Here, the responses of three cells in rat PGT08 are shown across all seven sessions in which they were present. It can be seen that in some sessions, these cells were robustly responsive to sucrose and in other sessions were completely unresponsive. In addition, within a session the vigor of the response was highly variable. This type of variability was apparent for all five tastants tested. **Fig. 2** shows the trial-to-trial and across-session variability in 11 cells recorded in rat PGT13. **Fig. 2A-B** show the normalized calcium signals (ΔF/F) recorded for portions of two sessions spaced one week apart. **Fig. 2C-D** show the corresponding evoked responses for all tastants tested. Note that cells C07 and CO8 did not respond to any taste stimulus. Most tastants evoked responses in the other cells across the one-week span. However, it is apparent that citric acid only evoked a weak or no response in several cells (C00, C01, C02, C04, C05, C09) in the first week (**Fig. 2C**) but evoked robust responses across all taste-responsive cells in the second week (**Fig. 2D**). Despite this trial-by-trial and across-session variability, across unit patterns associated with each of the tastants nevertheless remained discriminable. This is shown in **Fig. 3**. Here, the results of PCA analysis of recorded cells in rat PGT13 on day 8 and day 43 are shown. It can be seen that the responses to the five taste stimuli and artificial saliva remain separable in the PCA space. Thus, despite response variability within and across sessions, the ability of the population to identify and discriminate among taste qualities persists.

**Figure 2.**
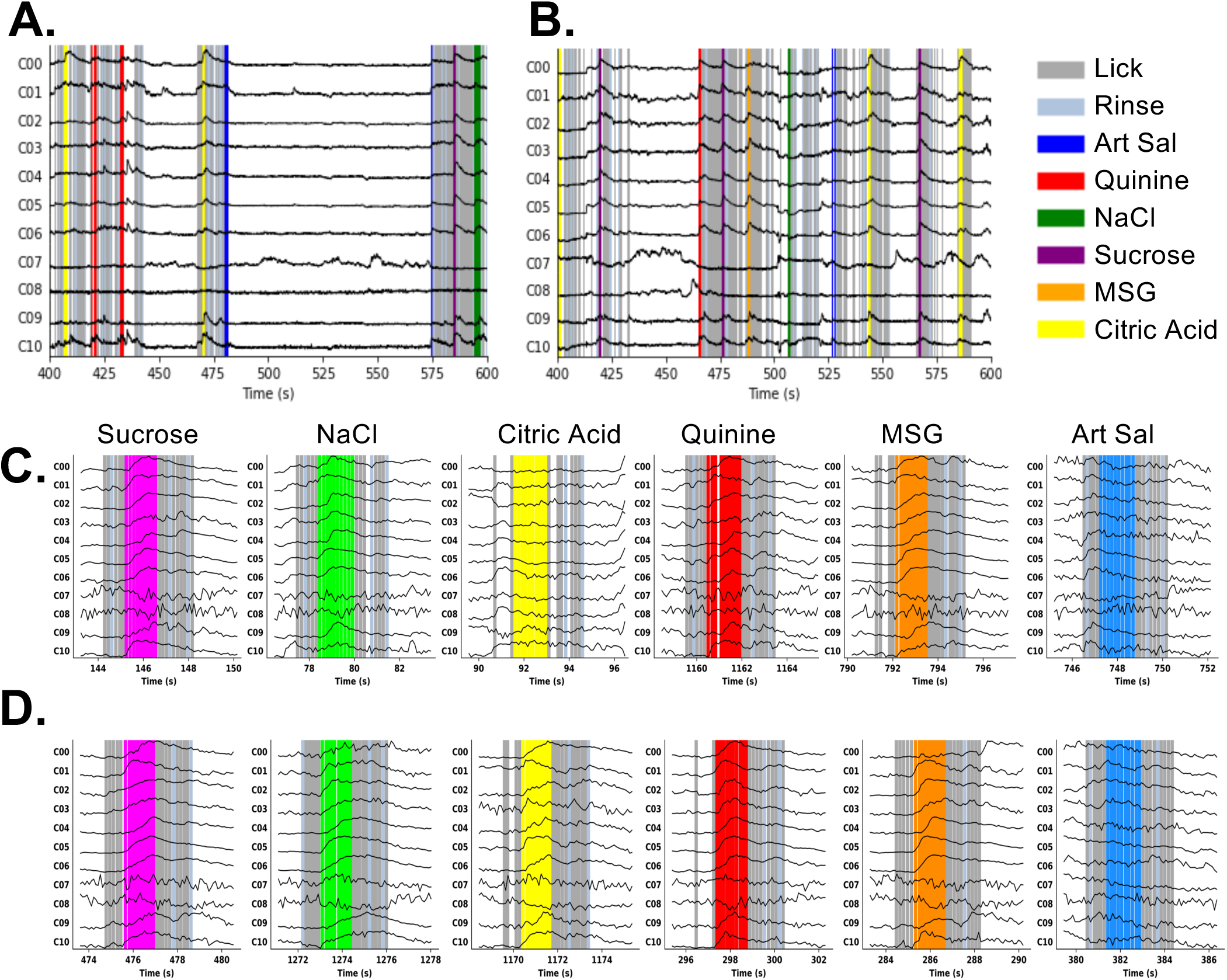
**A-B.** Examples of one-photon *in vivo* calcium imaging recording in two sessions, six days apart, from the same rat (PGT13). In this figure the term “rinse” refers to AS presented on a VR5 schedule. **C.-D.** examples of individual trials for each of the five tastants tested and AS. Examples in panel C. correspond to the session illustrated in A. and the examples in panel D. correspond to the session illustrated in B. The traces associated with each tastant across sessions show the variability in response to each of the tastants across cells. Each panel shows min–max normalized fluorescence traces. Normalization was performed independently per neuron and trial window by subtracting the minimum and scaling to the maximum within the window.

**Figure 3.**
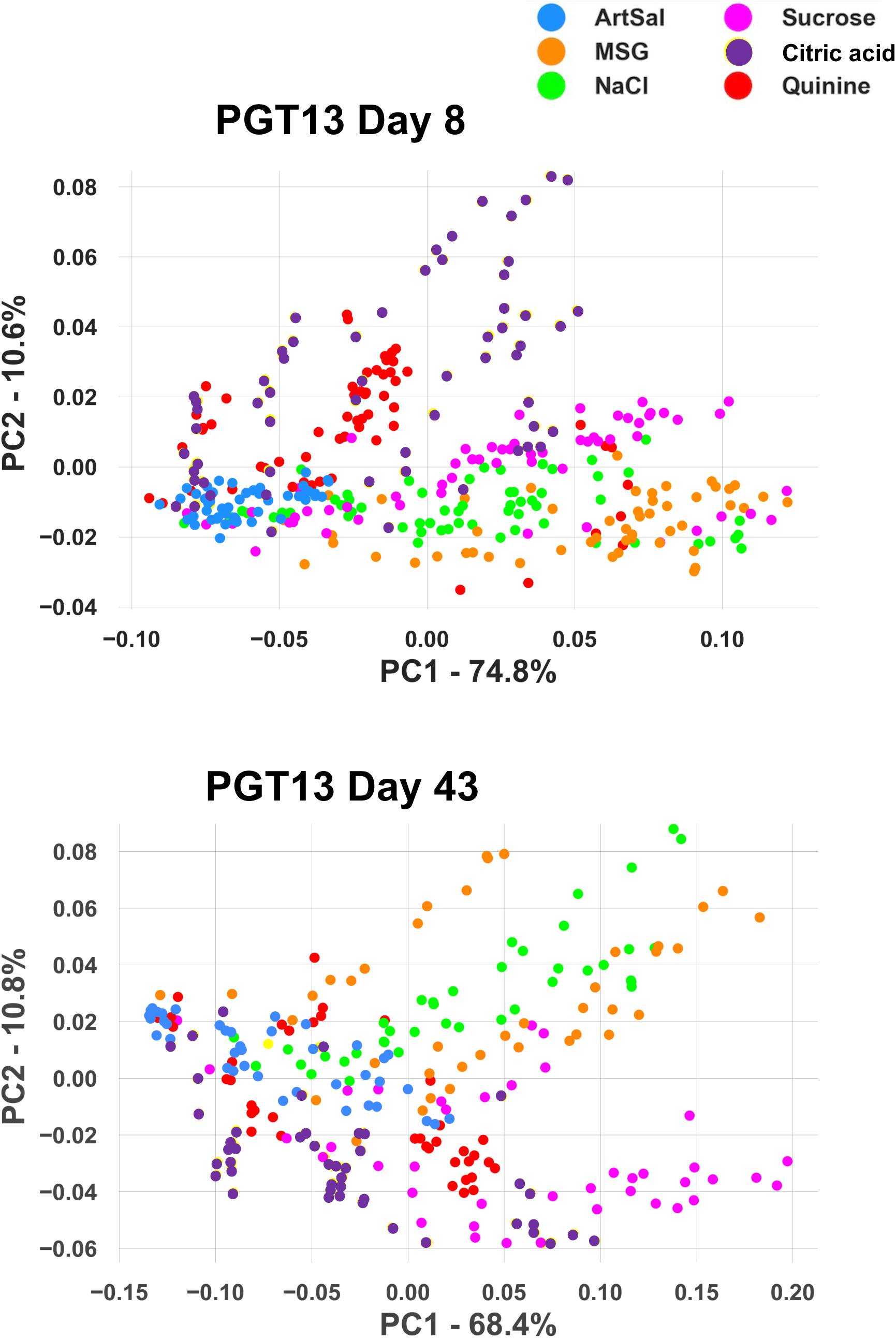
Results of PCA of taste responses imaged from the same 10 PbN cells in animal PGT13 on day eight (top) and day 43 (bottom). Each point represents responses to one lick of a stimulus. Even with only 10 cells showing variable response profiles across days, responses to the various taste qualities remained roughly segregated.

Changes in response magnitude resulted in significant changes in the breadth of tuning across sessions. **Figs. 4 and 5** show the changes in Uncertainty (*H*) across sessions in two animals, PGT08 and PGT13. In **Figs. 4A, 5A and 5D**, the Uncertainty measures across sessions were fairly similar. This was true despite dramatic changes in the response magnitudes evoked by the various tastants. **Figs.4B-C, 5B-C and 5E-F** show examples of the changes in tuning (**Figs. 4B, 5B, 5E**) across sessions in three cells. **Figs. 4C, 5C and 5F** show the response magnitudes evoked by all tastants tested corresponding to **Figs. 4B, 5B and 5E** respectively. Responses to tastants that were present across every session are shown in red bars. In **Fig 4C**, only one taste stimulus, quinine, consistently evoked a response, though it was not always the stimulus that evoked the best response across the tastants tested. In the two cells whose responses are depicted in **Figs. 5C** and **E**, there were several tastants that evoked responses consistently across sessions, though again, not always with consistent response magnitude. In contrast to the cells shown in **Figs. 4A-C and Fig. 5A-F**, the changes in tuning across sessions for cells in **Fig. 4D-F** showed relatively greater variability in the breadth of tuning across sessions, e.g., going from broadly tuned to narrowly tuned and back to broadly tuned, etc. The tuning changes (**Fig 4E**) and associated responses to tastants (**Fig. 4F**) in one cell show high variability across sessions. In this cell, there were no taste stimuli that evoked a significant response across all sessions. In several sessions, the cell appeared narrowly tuned to a single taste stimulus, though the identity of that stimulus was not consistent across sessions. In sum, despite the variability in response magnitudes within cells and across sessions, the breadth of tuning remained fairly stable for most cells. This may underlie the ability of the population to discriminate among taste qualities.

**Figure 4.**
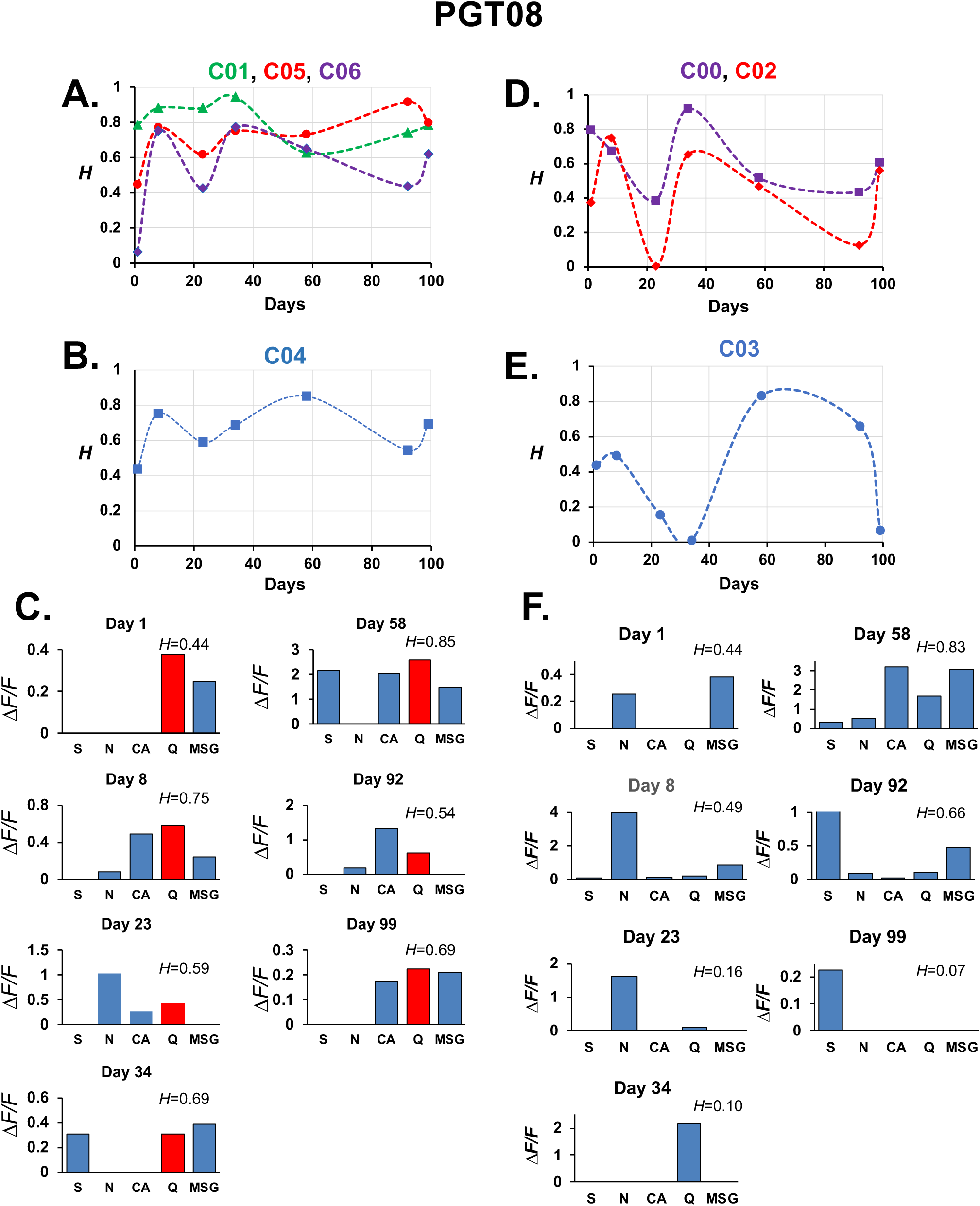
Response profile variability across days in animal PGT08. **A.-B.** changes in the Uncertainty measure of breadth of tuning (*H*) in four cells with relatively stable values of *H.* **C.** Responses to all tastants in cell C04 in all sessions. Red bars in each graph denote a stimulus that always evoked a significant response, though not always the best response compared to responses to other tastants. **D.-E.** Changes in *H* in three cells with widely variable values of *H* across days. **F.** Responses to all tastants in cell C03 in all sessions. In this cell, there were no taste stimuli that consistently evoked a response across all sessions recorded. Abbreviations: S, sucrose; N, NaCl; CA, citric acid; Q, quinine; MSG, monosodium glutamate.

**Figure 5.**
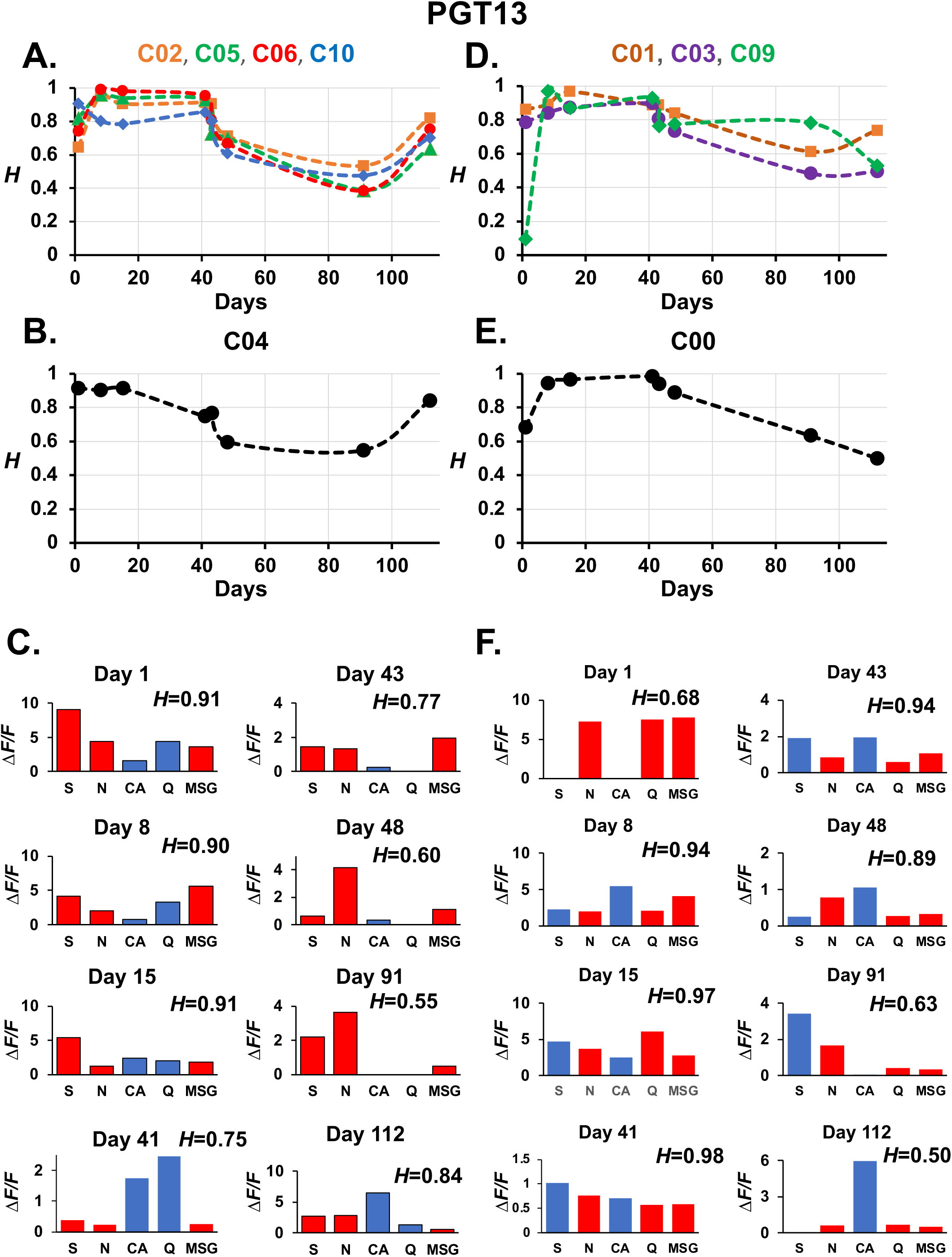
Response profile variability across days in animal PGT13. **A.-B.** changes in the Uncertainty measure of breadth of tuning (*H*) in five cells with similar values of *H* across sessions. **C.** Responses to all tastants in cell C04 in all sessions. **D.-E.** Changes in *H* in four cells across days. **F.** Responses to all tastants in cell C00 in all sessions. In C. and F., red bars in each graph denote a stimulus that always evoked a significant response, though not always the best response compared to responses to other tastants. Abbreviations: S, sucrose; N, NaCl; CA, citric acid; Q, quine; MSG, monosodium glutamate.

### Population coding of taste, olfaction and food

The ability to image the activity of a number of cells in the PbN simultaneously as they responded to various stimuli enabled the observation of, and subsequent theorizing about, population coding during appetitive and consummatory behaviors. Several prominent features of the PbN responses to tastants were apparent. For instance, as in the responses to sucrose and NaCl shown in **Fig. 6A**, responses to taste stimuli evolve gradually, recruiting more and more cells as the response unfolds over time. Of note is that, even though most cells respond to both sucrose and NaCl by the end of the response interval, the spatial and temporal sequence of responding cells differed between the two stimuli. As a result, even though most taste-responsive cells responded to both sucrose and NaCl (**Fig. 6B**), their spatiotemporal patterns of response were unique. In **Fig. 6B, left**, cells that responded to sucrose but not NaCl are shown as red-filled circles and in **Fig. 6B, right**, cells that responded to NaCl but not sucrose are shown as blue-filled circles. Thus, there are some cells that appear to be taste-selective.

**Figure 6.**
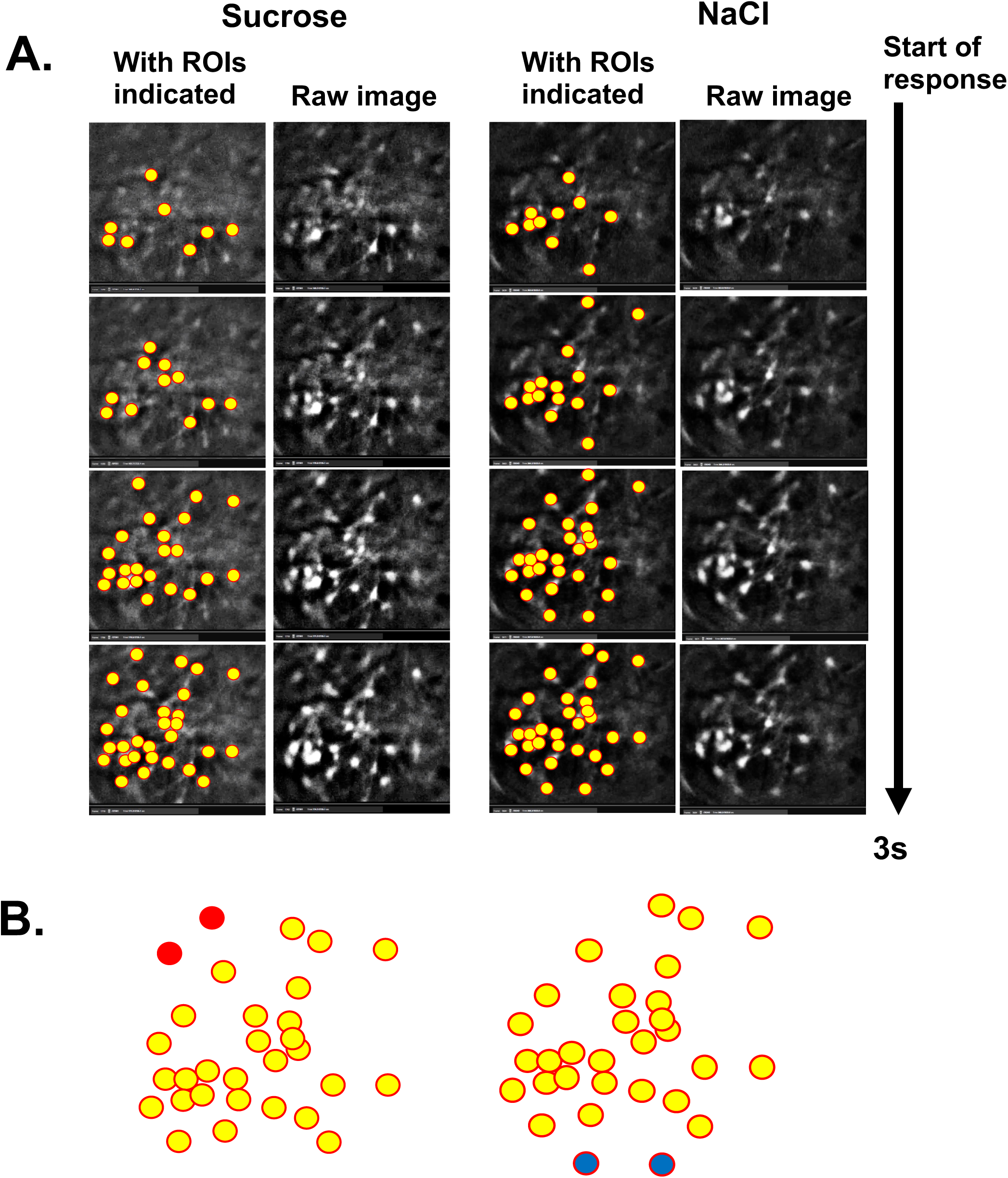
**A.** snapshots of responses to sucrose and NaCl across the first 3 s of response in animal PGT13. To the left of each raw image is the same image with regions of interest (ROI), i.e., cells, indicated by yellow-filled circles. For both sucrose and NaCl, more cells appear in the response as the response unfolds over time. **B**. Diagram of cells that responded to sucrose (left) and NaCl (right). Yellow-filled circles indicate cells that responded to both taste stimuli. Red-filled circles indicate cells that responded to sucrose but not NaCl, and blue-filled circles indicate cells that responded to NaCl but not sucrose.

Data show that licking evoked a shift in network activity that is sudden and distinct from that associated with other behaviors, e.g., grooming. **Fig. 7** shows the normalized activity of 11 cells recorded in one session from animal PGT06; **Fig. 7B** shows this activity from a selected portion of the session where the animal was licking taste stimuli. Taste responses are apparent in cells 2 and 3, while cells 7 and 8 became more active when the animal was not licking. These cells may be anti-lick cells [8, 9]. **Fig. 7C** is a snapshot of the PbN showing units that were present during the recording. Not every responsive cell is shown in this panel since some cells are not active until the appropriate stimulus is presented. **Fig 7D** shows the sequence of population network activity in a PCA space, with activity in sequential 100 ms bins joined by a line color-coded by the activity in which the animal is engaged. It can be seen that when animal begans to lick, the population activity was abruptly segregated from the activity generated when the animal was not licking. Examples of activity from two licking bouts are shown. **Fig. 8** is an analogous figure showing recordings from 15 cells in animal PGT08. In this case, taste-responsive, lick-related and anti-lick cells are indicated. These response types have been noted previously in electrophysiological experiments [8, 13]. **Fig. 8D** shows the results of a PCA analysis where population activity during licking, grooming, and other behaviors is shown to be well segregated. Notably, activity generated by licking is different from that generated by grooming, even though the two behaviors share a good deal of topography. Collectively, **Figs. 7D** and **8D** underscore the idea that appetitive behavior (licking) generates a unique pattern of neural population activity in the PbN.

**Figure 7.**
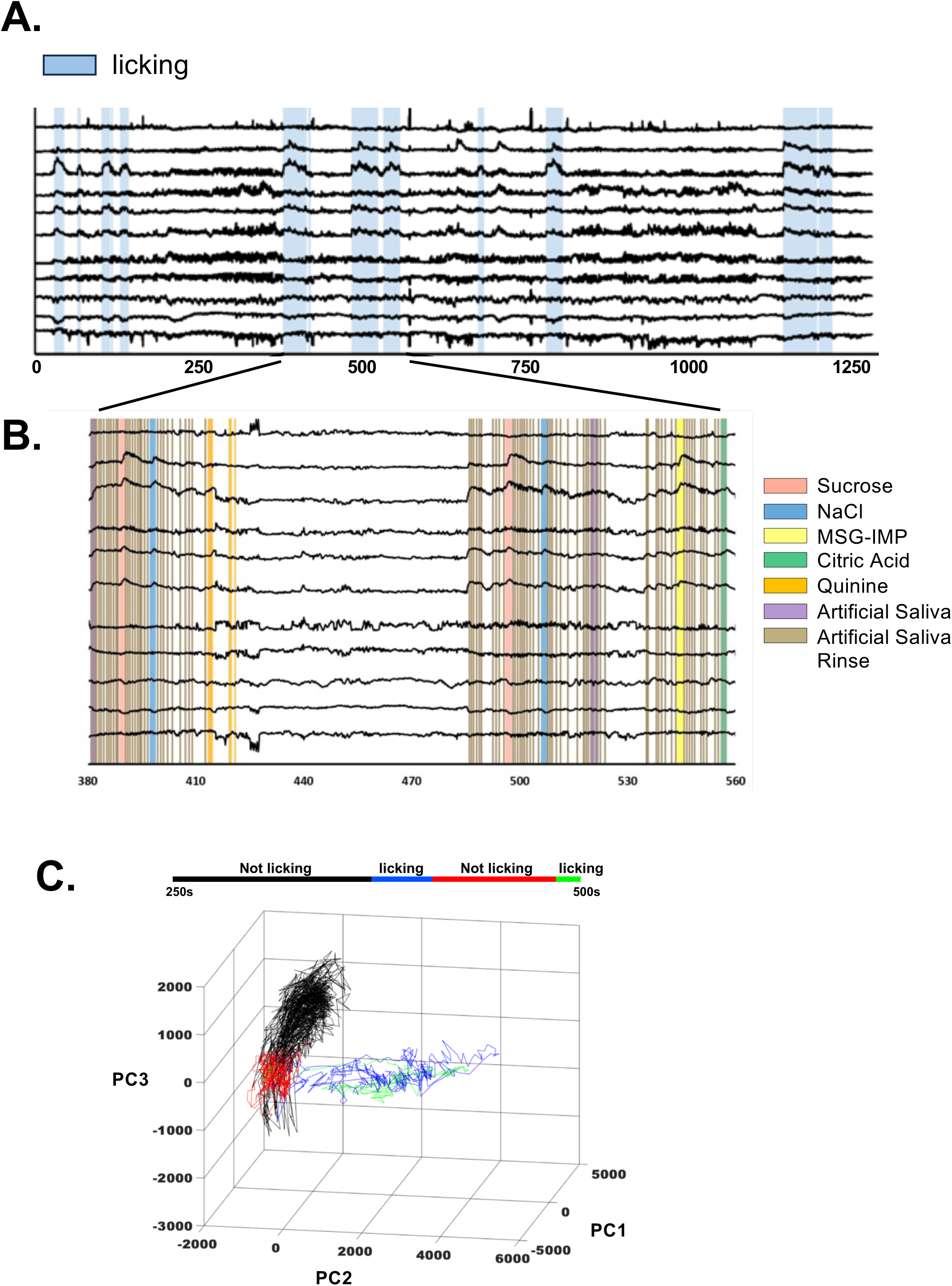
Results of calcium imaging in animal PGT06 during the Lick phase. **A.** Traces of calcium signals from 11 cells. Blue-shaded areas show intervals when the animal was licking taste stimuli. **B.** Magnification of calcium signal traces showing occurrences of specific taste stimuli as the animal licked. **C.** PCA analysis of activity during the Lick phase. Periods of licking generated a distinct pattern of network activity compared with periods when the animal was not licking.

Appetitive and consummatory behaviors associated with solid foods also evoked unique patterns of PbN population responses. **Fig. 9A, left**, shows the calcium responses while the animal was approaching the food well containing apple and subsequently eating its contents, as well as when the animal was grooming. **Fig. 9A, right**, shows the results of PCA analysis of this session. Here, it is apparent that, as in **Fig. 8D**, grooming evoked a different population response than that to food approach and eating. In addition, population responses to approach and eating were intermingled, suggesting that they may share a common trigger.

**Figure 8.**
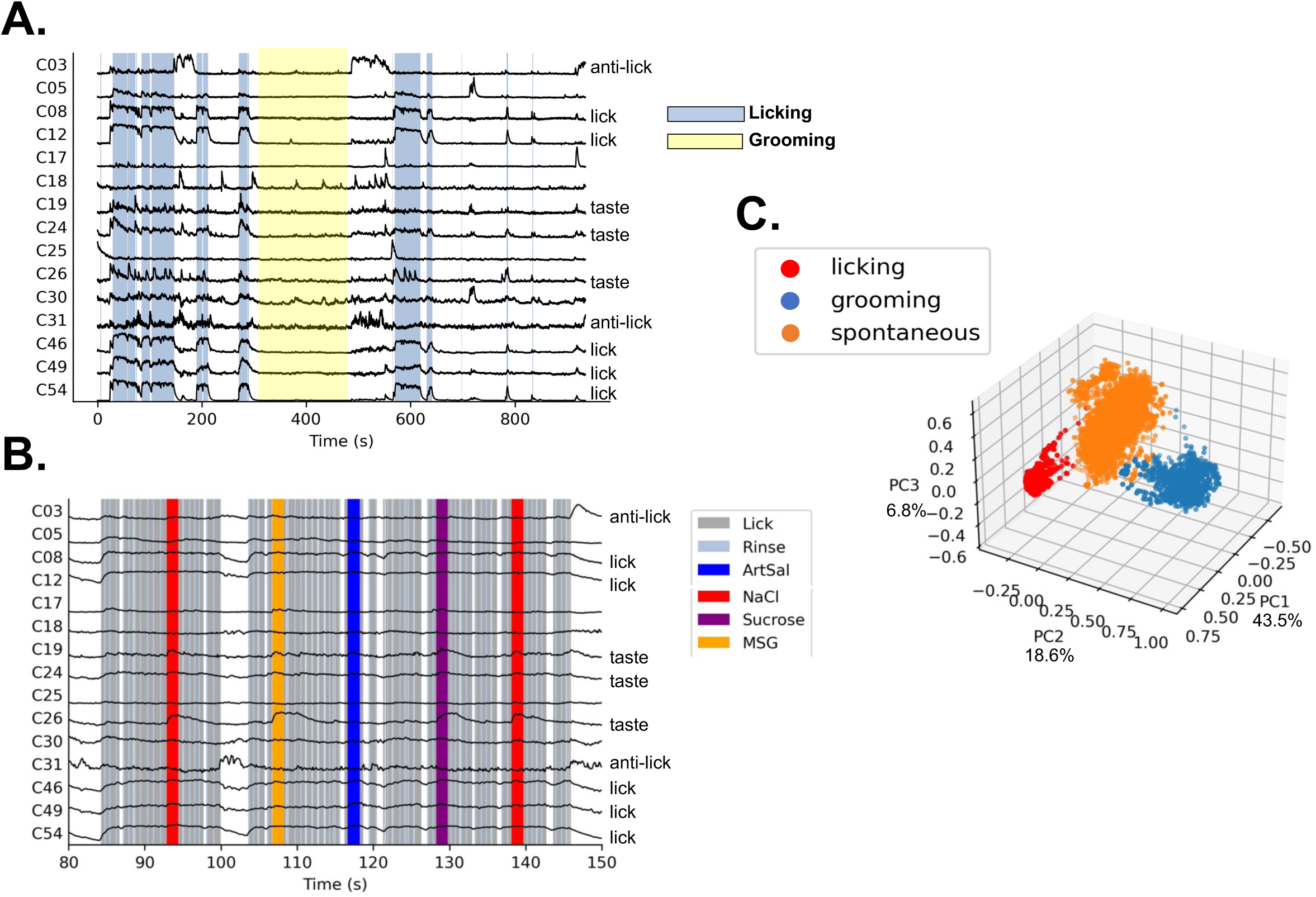
Results of calcium imaging in animal PGT08 during the Lick phase. **A.** traces of calcium signals from 15 cells. Blue-shaded areas show intervals when the animal was licking taste stimuli; yellow-shaded areas indicate intervals when the animal was grooming. **B.** Magnification of calcium signal traces showing occurrences of specific taste stimuli as the animal licked. **C.** PCA analysis of activity during the Lick phase. Periods of licking generated a distinct pattern of network activity compared with periods when the animal was not licking (spontaneous) or grooming.

**Figure 9.**
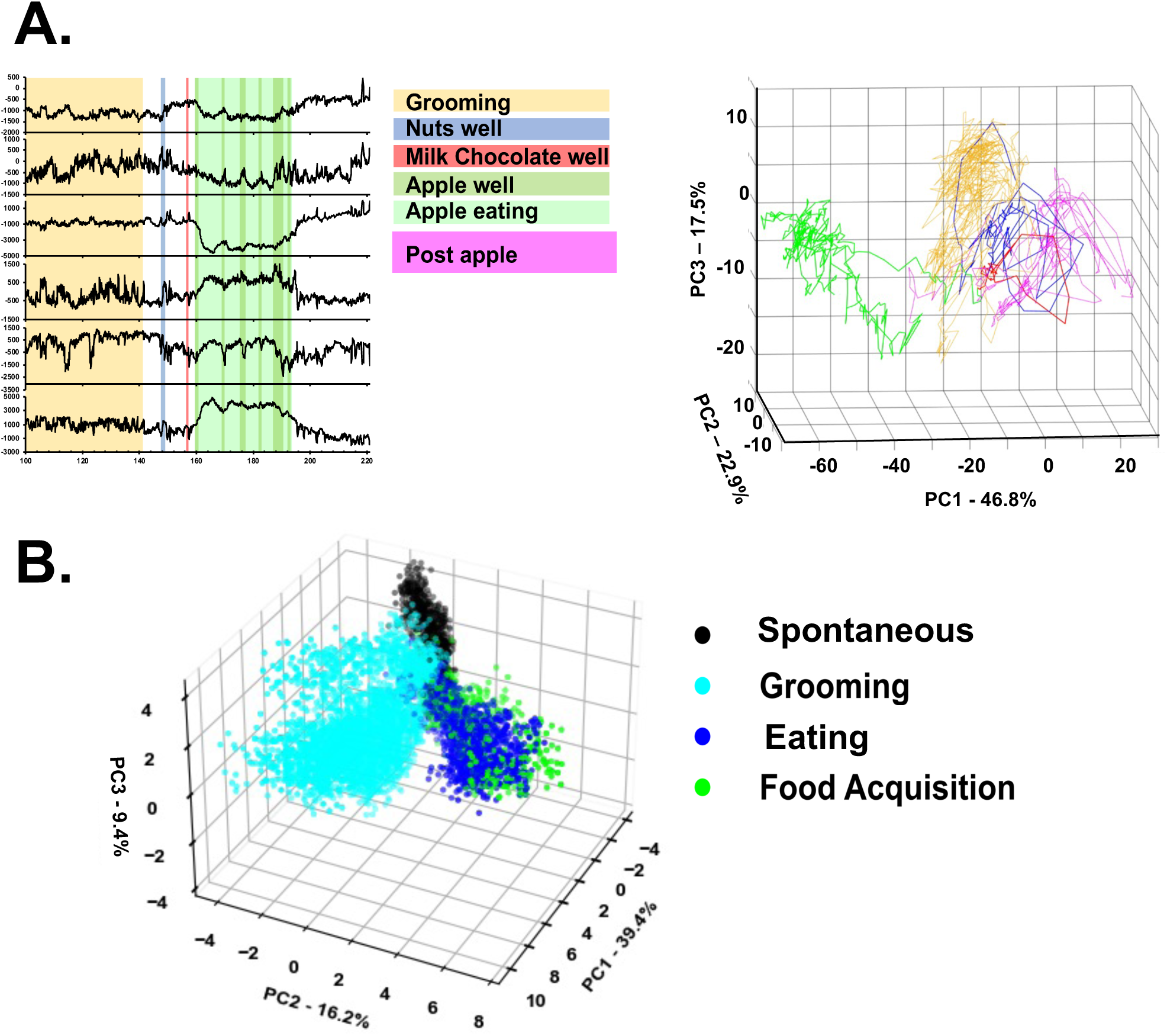
Results of calcium imaging in animal PGT09 during the Food phase. **A.** traces of calcium signals from 6 cells while the animal was either grooming (yellow-shaded area) or engaged in appetitive and consummatory behavior as indicated. Right, PCA analysis of activity during the Food phase shown on the left. Eating apple generated a pattern of network activity that was separate from that of all other activities. **B.** PCA of the entire session in the Food phase. Periods of eating and food acquisition generated a distinct pattern of network activity compared with periods when the animal was not eating (spontaneous) or grooming.

When an animal approaches the food well, part of the response may be due to olfactory stimulation. To test the feasibility of this idea, peanut and chocolate odors were presented in the Lick Phase portion of the experimental session in one animal. Results from 28 cells are presented in **Fig. 10**. Cells are arranged in order of response magnitude to sucrose. Several features of their responses are noteworthy. First, responses to citric acid, quinine, and both odorants are delayed past the point where stimulus licks end (see Fig. 1A). For citric acid and quinine, long latency responses in the PBN are not uncommon [8]. Four both peanut and chocolate odors, robust responses occurred after the AS licks ended. Second, although one might expect that the same cells that responded to sucrose and NaCl would also respond to chocolate odor and peanut odor, respectively, this was not entirely the case. **Fig. 11A** shows the cells that responded to NaCl, sucrose, peanut odor and chocolate odor, as well as the cells that responded when the animal ate peanut or chocolate. As shown in **Fig. 11B-C**, for each stimulus comparison, there are cells that responded to both stimuli and others that responded to one or the other stimulus but not both. Similarly, as shown in **Fig. 11D**, some cells responded to eating either chocolate or peanuts and others that responded selectively to one food or the other. Collectively, these data show that PbN cells respond to the odor of food as well as the consumption of food. Moreover, responses to liquid tastants, solid foods and their odors each include cells that are selective to that stimulus.

**Figure 10.**
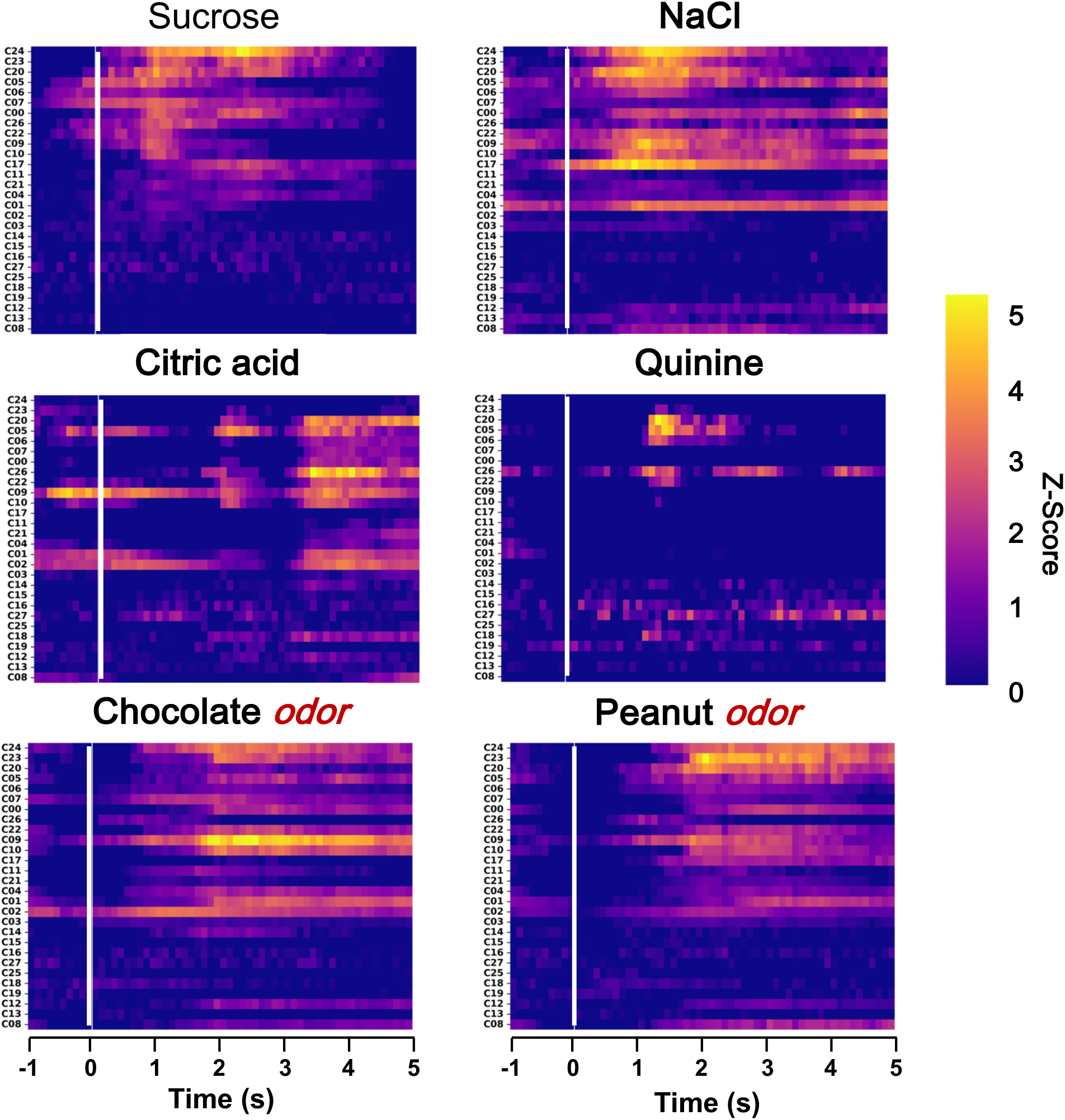
Heat maps of normalized, calcium signals from 28 cells in animal PGT 13. Cells are arranged from top to bottom according to the magnitude of the response to sucrose. Six seconds of activity are shown for each stimulus: 1 s baseline and 5 s of response. Vertical white line indicates the beginning of the stimulus presentation. Both taste stimulate and food odors evoke robust responses across overlapping sets of cells. Note that responses to odorants

**Figure 11.**
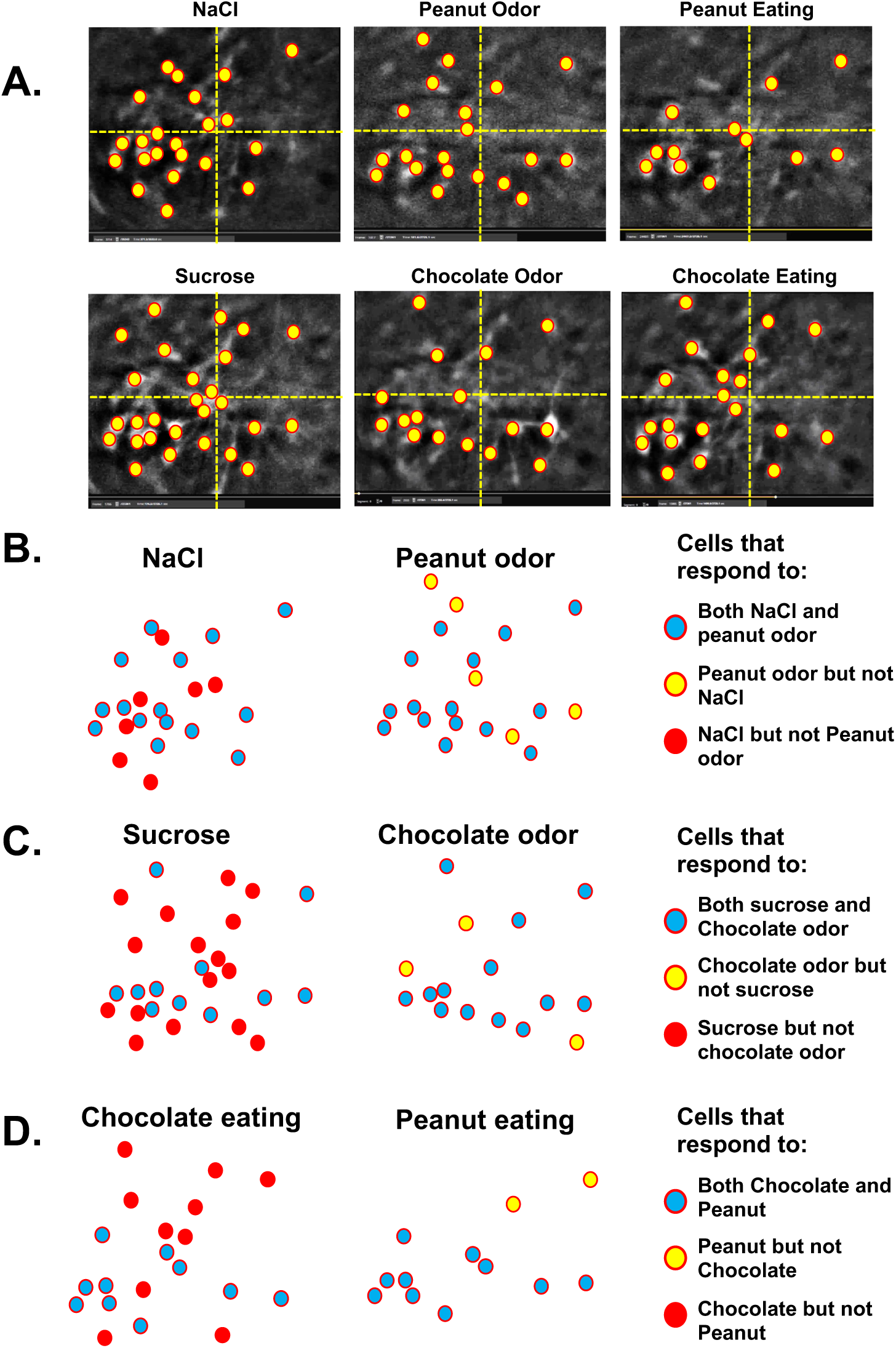
Comparison of PbN responses to NaCl taste, food, odors, and eating. **A.** Snapshots of the PbN showing the cellular responses to NaCl, peanut odor, peanut eating, sucrose, chocolate, odor, and chocolate eating. Yellow-filled circles indicate active cells. Vertical and horizontal dashed lines divide the field into quadrants. **B.-D.** Diagrams of responsive cells showing the overlap across stimuli. For each comparison, there are some cells that respond to one stimulus, but not the other.

Population coding of appetitive and consummatory behaviors differs from that associated with licking various tastants. Activity associated with both of these behaviors differs from that evoked by grooming. **Fig. 12A** shows the calcium-evoked traces from 28 cells for an entire session where the animal licked taste stimuli and ate solid foods. Close examination of this figure reveals the heterogeneity of cellular responses in this sample of cells. For example, although most cells became more active during licking and grooming, a smaller subset of cells were particularly active during eating. In addition, there are cells such as C19 and C25, which appeared to be more active between lick bouts than during lick bouts. These cells might be classified as anti-lick cells [8, 9]. **Fig. 12B** depicts a summary of population activity during the entire session using PCA analysis. Importantly, activity associated with licking tastants was nearly orthogonal to that generated by appetitive and consummatory behaviors associated with solid foods. Activity during tastant licking was roughly, though imperfectly, segregated by taste quality. Odor-evoked activity was most closely related to tastant-evoked activity rather than eating food.

**Figure 12.**
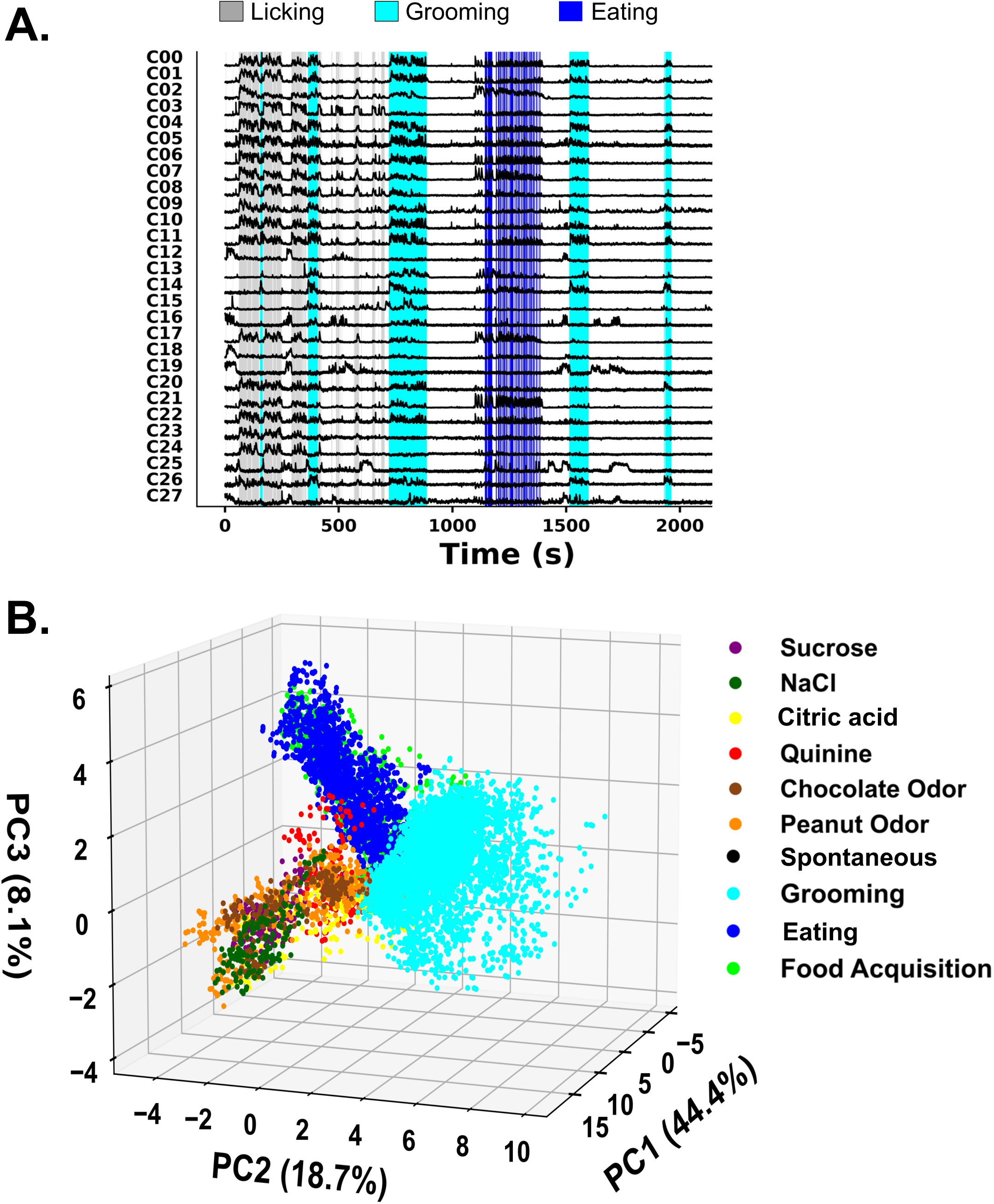
Summary of calcium imaging results in the PbN of animal PGT13 during the Licking and Food phases. **A.** Calcium traces from 28 cells in the PbN as the animal licked, groomed, and approached and ate food. **B.** PCA of network activity generated by 28 PBN cells as the animal licked tastants, groomed, and approached and ate food. Responses to licking taste stimuli and sniffing food odors were segregated from the network activity generated by acquiring and eating food. Activity generated by approach to the food well is hidden by the activity generated by eating in this figure. The network activity generated by these two behaviors was segregated from that evoked by grooming, despite the overlap in motor topography.

## Discussion

In the present study, we used one-photon calcium imaging to investigate two aspects of the neural representation of taste and food in the PbN in awake, unrestrained rats: tuning profiles of individual cells and the characterization of responses to tastants and solid food across the network. The technical demands of accessing this small, deep nucleus limited the number of cells that could be recorded. Nevertheless, this study provided a window on how the activity patterns of these neurons related to their tuning properties determined from standard liquid tastants, how their activity was modulated by naturalistic eating and other behaviors, and how tuning properties varied over time. Analyses showed that PbN cells were generally broadly tuned across taste qualities, in agreement with previous electrophysiological [8, 13] and calcium-imaging [14] studies. However, their tuning profiles and breadth of tuning changed, sometimes radically, across days/weeks. Despite this wide-ranging variability, the across unit patterns of activity could nevertheless distinguish among the basic taste qualities. Responses to a given tastant began in a small group of cells but quickly expanded to incorporate more and more cells over time. This gave the responses to tastants unique spatiotemporal patterns. Shifts in population network activity notably differentiated activity related to licking tastants, to appetitive or consummatory behavior and to activity related to grooming. Presumably, these distinctive network patterns reflected both motor activity and the sensory properties of tastants and food. Importantly, we found that PbN cells responded to food odors in addition to responses to tastants and solid food. Although there was much overlap in the cells that responded to taste stimuli, to the odor of a food and to eating that food, there were cells that were very specific, though these were in the minority. In all, present data suggest that taste-responsive neurons in the PbN evidence changes in their tuning profiles over time, while the representation of taste qualities remains discriminable through population activity. Further, in addition to responses to taste quality, neurons in the PbN also respond to food-related odors and have activity patterns that are modulated by appetitive and consummatory behaviors, underscoring the sensorimotor function of the PbN.

### Taste response properties

Data reported here show that taste-responsive cells in the PbN express broad tuning across taste qualities. Imaging results from PbN cells reported here agree with previous electrophysiological studies in awake rats [8, 13] in that these cells generally responded to more than one taste quality. Moreover, Jarvie et al. [14], using one-photon calcium imaging in mice, showed that taste-responsive neurons in the PbN expressing the Satb2 gene generally respond to multiple basic taste qualities. To incorporate observations of single taste cells responsive to more than one taste quality, the labeled line theory argues that each cell responds “best” to one tastant, and that a cell’s best stimulus defines its function in the neural code for taste (reviewed in [7]).

Perhaps an unwritten assumption of the labeled line theory is that the response properties of best-stimulus cell types are stable features of the cells, such that a sucrose-best cell will always be a sucrose-best cell, a salt-best cell will always be a salt-best cell, and so on (see [15]). However, there are several studies, including the present one, suggesting that this assumption may be false. For example, Shimatani et al. [16] showed that the response profiles of chorda tympani fibers, innervating taste buds on the rostral tongue, changed over days. Moreover, by recording from the same cell over multiple days, Sammons et al. [17] found that cells in the NTS and PbN showed similar changes. To account for these results, Shimatani et al. [16] and Sammons et al. [17] suggested that changing input to peripheral taste nerve fibers might be at play. It is well known that there is frequent turnover of the taste receptor cells in tastebuds on the tongue [18]. The reinnervation of the freshly born taste receptor cells, may be the source of variable response properties of the cells in the taste buds [19]. Collectively, these data point to the idea that, on a cell-by-cell basis, response properties of taste cells in the brainstem are changeable, casting doubt on a strict labeled line conceptualization of taste coding.

The type of response profile variability that was observed in PbN cells in the present study is reminiscent of what has been called “representational shift” by Schoonover et al. [20] in the olfactory system. This phenomenon has also been reported in other sensory systems including somatosensation [21], vision [22], audition [23] and gustation [17]. In the present study, where we could observe multiple cells simultaneously, the critical question is what aspect of the global representation of taste is stable such that the upstream “readers” of the signal can identify and discriminate among taste stimuli. In the case of the PbN, despite the relatively small sample of the population that was imaged, PCA analyses showed that the across unit patterns evoked by the various tastants were separable in the “taste space”. This was apparent, even when the PCA analyses were conducted weeks apart.

### PbN network responses to taste and food

The ability to image multiple PbN cells simultaneously permitted the observation of a temporal evolution of taste responses as the animal licked various tastes. Specifically, differing latencies of response across the sample of responsive cells produced unique spatiotemporal patterns of response. Beginning with the first lick of a taste stimulus, just a few cells began to respond with more and more cells joining the response over the next couple of seconds. It is possible that this result may be an artifact of the calcium dynamics of different cells, or the position of the cells with respect to the GRIN lens. However, if the differences in the temporal evolution of the responses to sucrose and NaCl were influenced by either of these factors, one might expect that the sequence of active cells would be similar, which was not the case (see Fig. 6.). In agreement with present results, differences in the latency of responses in the PbN have been documented in experiments in awake unrestrained animals [8], albeit from electrophysiological recordings of single cells recorded one at a time. The dynamics of the unfolding of each taste response over time producing unique spatiotemporal patterns of activity may be another way of conveying information upstream.

In addition to responses to the basic taste qualities, we also recorded responses to food odors, i.e., peanut and chocolate odors, in PbN cells. Electrophysiological responses to odorants in both the NTS [24, 25] and PbN [26] have been reported previously. So, it is not surprising that these responses were evident in the present data. However, the origin of these responses remains a mystery. Responses to odorants in both the NTS and PbN are likely the result of top down activity since there are no known direct projections from olfactory-related areas to either structure. However, it is unknown whether the olfactory responses in the PbN are the result of input from the NTS or whether the PbN receives an olfactory-related input independent of that to the NTS. It is notable that the responses to each of these two food odors were not identical to the responses to eating the corresponding foods, though there was much overlap. It is possible that the origins of these two responses may not be identical. For example, responses to odorants may reflect centrifugal influence, either relayed through the NTS or directly to the PbN, while responses to foods likely originate in the periphery.

In apparent contrast to previous work in the NTS [10], there was no consistent increase in cellular activity when the animal’s head was inside the food well, and no widespread suppression of activity when the animal was eating. It is difficult to draw conclusions from these apparently contrasting observations in the NTS versus the PbN because the techniques for observing cellular activity were so different. That is, an inhibitory response in the PbN might be noted by the absence of cellular activity, while in the NTS an attenuation of firing rate could be observed. When the animal was eating peanuts, only about a third of the cells in the NTS were excited and 2/3 were suppressed [10]. In the PbN, there were fewer cells that were active compared to when the animal was sniffing peanut odor (∼60%) or licking NaCl (∼55%). Trial to trial variability make these numbers approximate. The relative paucity of cells that were excited when the rat was eating peanuts might reflect the inhibition of other cells that were part of the response, but not excited. In the case of chocolate, there appeared to be about the same number of cells responsive to the odor of chocolate and the eating of chocolate, but fewer than the number of cells responsive to licking sucrose. In all, differences in the methodology used to study the NTS vs. the PbN responses to food make direct comparisons problematical, though there are some data that point to consistent reactions.

Examination of network activity, as characterized by PCA analyses, in two animals that were permitted to eat solid food revealed differences in both motor- and sensory-generated activity in the PbN. For example, PCA analyses in both animals while they approached the food well, ate food or groomed, showed that approach and consumption of food generated similar network activity that was clearly different from that generated by grooming, despite the similarity in motor typography. In one animal that was also allowed to lick tastants and smell food-related odors, network activity generated by licking and sniffing activity was clearly different than, and nearly orthogonal to that generated by approach to and consumption of food, or grooming (see **Fig. 12**). In the portion of the PCA results associated with licking tastants and smelling food odors, there was a rough segregation of activity associated with each tastant and odorant. The overall segregation of network activity associated with taste and licking and smelling odorants versus approach and eating solid foods versus grooming, may be the result of different movements supporting each of these behavior categories. For example, responses to odorants occurred during dry licks, perhaps accounting for their close association with responses to liquid tastants. Since there are no known direct or indirect motor pathways that project to the PbN, the most plausible source of motor-related activity in this area may be somatosensory feedback generated by the movements. Even so, the fact remains that cellular activity in the PbN related to taste and food is sensorimotor rather than strictly sensory in nature.

### Caveats and limitations

The present study is not without its caveats, some of which may limit the conclusions to be drawn. First, the recording of calcium signals is not equivalent to spiking. So, although it is tempting to suggest a one-to-one relationship between the relative strength of the calcium signal and the number of spikes produced by the neuron, it is only speculative at this point. Thus, differences in response latency may be a reflection of differences in the strength of the calcium signal, and the ability of our apparatus to detect it. Nevertheless, differences in response latency across stimulate can be informative. Second, it is impossible to identify whether an active neuron is inhibitory or excitatory, which limits the conclusions about neural circuitry that can be drawn. Third, the position of the GRIN lens with respect to the responsive cells naturally limits how much of the PbN can actually be captured. We would argue, however, that the conclusions we have drawn about the cells that we have imaged would not be affected by imaging more cells. That is, even with the limited number of cells that were imaged, the phenomena that we describe are present but how widespread they are is unknown. Fourth, our sample of cells is relatively small; however, the advantage of imaging a group of cells that are responding to taste and food in the same animal at the same time has revealed important features of the PbN that could not have been obtained by other means.

### Conclusions

The PbN has traditionally been characterized as a “taste relay,” serving as an obligatory synapse in the central gustatory pathway of non-primate mammals. The present study, however, has added an expanded perspective to the functionality of the PbN. By imaging multiple PbN cells simultaneously and sometimes over long periods of time, it was possible to discover several new features of PbN responses to tastants and solid food. For example, by observing the pattern of recruitment of cells during a response to taste, unique spatiotemporal patterns associated with various tastants could be observed. Importantly, data showed that tuning profiles of individual cells changed over time periods of days to weeks; however, the across unit patterns of response to the various taste qualities remained discriminable. In addition, PbN cells responded to food odors, and to the consumption of solid food. However, the patterns of population activity associated with responses to food odors were more closely associated with those to licking taste stimuli than they were to eating the corresponding food. Further, information about taste, olfaction and motor-related somatosensory input appear to be expressed in overlapping sets of PbN cells, rather than in segregated groups. Collectively, these data broaden the response repertoire of the PbN to include sensorimotor functions related to appetitive and consummatory behavior, as well as taste-related neural representation.

## Author contributions

Conceptualization, P.D.; Methodology, P.D., J.S., and F.O.; Software, F.O., S.P.; Formal Analysis, F.O., S.P.; Investigation, F.O.; Data Curation, F.O.; Writing – Original Draft Preparation, P.D.; Supervision, P.M.; Project Administration, P.D., F.O.; Funding Acquisition, P.D.

## Institutional Review Board

The study was conducted according to the guidelines of the Declaration of Helsinki, and approved by the Institutional Animal Care and Use Committee of Binghamton University (Protocol 846-21, *Temporal Coding in the Gustatory System of the Brain*, approved February 25, 2021).

## Data availability

https://doi.org/10.6084/m9.figshare.29963645.v1

## Acknowledgements

The authors wish to thank Dr. Jonathan D. Victor for valuable comments and suggestions related to this work.

